# An epiphytic tissue sampling method using a cellulose acetate membrane filter facilitates phylogenomic analysis of *Plasmopara viticola*

**DOI:** 10.64898/2026.11.13.765879

**Authors:** Chisako Kida, Shuji Shiozaki, Koh Aoki, Ayako Tsushima

**Affiliations:** Graduate School of Agriculture, Osaka Metropolitan University, Sakai, Japan

**Keywords:** cellulose acetate, grapevine downy mildew, peel technique, phylogenetic analysis, *Plasmopara viticola*, single nucleotide polymorphisms

## Abstract

Understanding genetic backgrounds and potential migration routes of emerging pathogens is important for preventing further invasions and outbreaks. RNA-seq data from field-collected infected plant tissues are increasingly used for pathogen monitoring, particularly for pathogens that cannot be cultured on artificial media. Despite its advantages, low pathogen read proportions in such field RNA-seq data can lead to unreliable genotyping results, hindering accurate surveillance and diagnosis. *Plasmopara viticola*, the causal agent of grapevine downy mildew, is one of the most destructive grapevine pathogens. Historical records and previous studies suggest that this pathogen originated in North America and has spread worldwide, including to Japan. However, the phylogenetic relationship between *P. viticola* populations in Japan and other geographic regions remains unknown, largely because culturing this obligate biotrophic pathogen is labor-intensive. Here, inspired by a peel technique used in paleobotany, we developed a new sampling method that effectively collects epiphytic *P. viticola* material using a cellulose acetate membrane filter. Samples collected using this method yielded higher RNA quantity and quality than conventional leaf discs. Moreover, the new method significantly enriched pathogen biomass relative to host tissues. By integrating newly obtained field RNA-seq data and publicly available whole-genome sequencing (WGS) data, we conducted single nucleotide polymorphism (SNP) calling and phylogenetic analysis. Our results indicated that the Japanese samples tested are genetically distinct from isolates mainly collected in European countries. In conclusion, this new sampling technique simplifies field sample collection of this agriculturally important plant pathogen and provides insights into developing better-informed management strategies.

## Introduction

Emerging and reemerging pathogens pose a serious threat to food security (Ristaino et al. 2021). Outbreaks of such pathogens are expanding and exacerbated by transmission through global transportation and trade networks, climate change, and/or evolution of new pathogen lineages (Inoue et al. 2017; Lewis et al. 2024; Bradshaw et al. 2025). Understanding the genetic background and estimating potential migration routes is key to controlling further invasion of these pathogens. To this end, researchers have performed population genetic analyses of plant pathogens using microsatellite markers that consider a limited number of genetic loci (Gross et al. 2014; Saleh et al. 2014). More recently, whole-genome sequencing (WGS) data generated by next-generation sequencing (NGS) technology enable non-targeted, comprehensive analysis of pathogen genomes. High-resolution single nucleotide polymorphism (SNP) profiles derived from WGS data are now widely used to reveal phylogenetic relationships, population structure, potential invasion routes, and genetic variations relevant to pathogenicity or fungicide resistance (Feurtey et al. 2023; Latorre et al. 2023).

However, WGS data cannot be readily applied to a subset of plant pathogens that cannot be cultured on artificial media. For DNA extraction and subsequent WGS data acquisition, purifying and propagating such obligate biotrophic pathogens on their hosts is tedious and time-consuming. To overcome this problem, RNA-seq data generated directly from pathogen-infected field plant samples have been used to monitor important obligate biotrophic pathogens, represented by *Puccinia* fungi causing wheat rust diseases (Hubbard et al. 2015). Using such field RNA-seq data, previous studies have successfully revealed population structures and potential migration routes of these pathogens by analyzing SNPs in expressed genes (van Schalkwyk et al. 2022; Tsushima et al. 2022). However, pathogen read proportions in field RNA-seq datasets are usually limited. For example, the average mapping rate to the pathogen reference genome was 48.2% for *P. striiformis* f. sp. *tritici* and 42.2% for *P. graminis* f. sp. *tritici* (van Schalkwyk et al. 2022; Tsushima et al. 2022). These restricted pathogen read proportions can reduce the number of detected SNPs and hinder reliable phylogenetic analyses. In addition, although a standard RNA extraction protocol yielded good results for *Puccinia* fungi-infected wheat leaves in previous studies, this is not always true for other plant tissues. For example, abundant polysaccharides and polyphenols in many fruit trees inhibit isolation of high-quality nucleic acids (Japelaghi et al. 2011). Thus, increasing the pathogen/host biomass ratio in field samples by eliminating plant tissues should improve RNA extraction and subsequent data analysis. This will, in turn, enhance plant pathogen surveillance and diagnosis using field RNA-seq data.

Grapevine downy mildew, caused by the obligate biotrophic oomycete *Plasmopara viticola*, is one of the most devastating diseases in viticulture worldwide (Bois et al. 2017). *P. viticola* is endemic to North America and was introduced to Europe in the 1870s, probably along with imported American *Vitis* species in attempts to identify phylloxera-resistant germplasm (Gessler et al. 2011). Fontaine et al. reported that the worldwide invasive *P. viticola* population originated from a native cryptic species (also called *forma specialis*), *P. viticola* f. sp. *aestivalis*, and that Europe served as a bridgehead for the spread of this pathogen (Fontaine et al. 2021). In Japan, *P. viticola* is suspected to have been introduced from overseas with infected nursery stocks (Aoki and Suzuki 2025; Yamada and Sato 2016). This hypothesis is based on historical records describing the first official report of *P. viticola* in Japan in 1890 (Tanaka 1890), following the initial introduction of grapevine seedlings from the United States in 1872-1873, and subsequently from France and China (Kikuchi 1948; Nagasawa 1976). However, the invasive route and genetic background of *P. viticola* in Japan have not been evaluated in detail. This is largely due to its obligate biotrophic nature, as isolating and culturing *P. viticola* requires tedious and time-consuming protocols.

The purpose of this study is to develop a technique to enrich *P. viticola* biomass in field-collected samples and to apply it to reveal genetic relationships among pathogen populations in Japan and other geographic areas. To this end, we have developed a sampling method for collecting epiphytic tissues where *P. viticola* forms sporangia, using a cellulose acetate membrane filter. We then conducted phylogenetic analyses of *P. viticola* from Japan and other countries using our original field RNA-seq datasets and publicly available genomic datasets.

## Materials and methods

### Plant materials

*P. viticola*-infected expanded grapevine leaves were collected at the Osaka Metropolitan University Research and Education Field in Sakai-shi and at a commercial farm field in Yamanashi-shi, Japan, from May to July in 2022 and 2023 (Supplementary Tables 1 and 2). Samples were collected between 9:30 a.m. and 10:00 a.m. Infected leaves were collected from seven grapevine cultivars (‘Aki Seedless’, ‘Campbell Early’, ‘Himrod’, ‘King Dela’, ‘Koshu’, ‘Kyoho’, and ‘Muscat Bailey A’) and *Vitis ficifolia* var. *ganebu*, which was originally collected in Uka, Okinawa, Japan, and propagated at the Osaka Metropolitan University Research and Education Field.

### Previously reported sampling methods using a cellulose acetate solution

We prepared a cellulose acetate solution (5% cellulose acetate dissolved in acetone) as described in Li et al. 2019. The cellulose acetate solution was applied to *P. viticola*-infected leaves of ‘Aki Seedless’ in three different ways, as explained in the results section.

### Epiphytic tissue sampling method using a cellulose acetate membrane filter

Infected leaves were detached by hand while wearing disposable nitrile gloves and placed in a resealable plastic bag. Peel samples using the newly established method were prepared as follows. A 13-mm-diameter cellulose acetate membrane filter (C045A013A; Advantec Toyo Kaisha, Ltd., Japan) was placed onto a lesion. Approximately 100 µL of acetone was applied to the membrane filter to attach it to the lesion site. After the membrane filter adhered to the leaf surface (which takes about three to five minutes, depending on humidity), the fixed membrane filter was removed from the leaf surface using forceps. For comparison, leaf disc samples were prepared by excising the lesion sites using a 12-mm-diameter cork borer. Each sample prepared using the two methods was placed in a sterilized 2.0-mL tube and snap-frozen in liquid nitrogen. When liquid nitrogen was unavailable, the sample was immersed in RNA*later* Stabilization Solution (Thermo Fisher Scientific, Inc., MA, USA) in a sterilized 2.0-mL tube to maintain nucleic acid integrity. After returning to the laboratory, the RNA*later* Stabilization Solution was removed from the tube by pipetting, and the sample was snap-frozen in liquid nitrogen. The snap-frozen samples were stored at −80°C until RNA extraction. To observe *P. viticola* embedded in peel samples, the samples were stained in 0.5% trypan blue solution (Nacalai Tesque, Inc., Japan) for 1 h at room temperature and washed twice with distilled water. The stained samples were observed using BX53 microscope (Olympus Corporation, Japan).

### RNA extraction and RNA-seq data generation

The samples were homogenized with a sterilized 5-mm-diameter metal bead using TissueLyser LT (QIAGEN N.V., The Netherlands) at 50 oscillations/second for 2 min. The homogenized samples were subjected to total RNA extraction using RNeasy Plant Mini Kit (QIAGEN N.V.), followed by DNase treatment using TURBO DNA-*free* Kit (Thermo Fisher Scientific, Inc.). The quantity and quality of RNA were assessed using NanoDrop Lite (Thermo Fisher Scientific, Inc.), Qubit RNA HS Assay Kit (Thermo Fisher Scientific, Inc.), and 4200 TapeStation system (Agilent Technologies, Inc., CA, USA). Samples subjected to RNA-seq are shown in Supplementary Tables 1 and 2. Genome-Lead Co., Ltd. (Japan) constructed complementary DNA libraries using KAPA mRNA Capture kit (Roche Diagnostics GmbH, Germany) and MGIEasy RNA Directional Library Prep Set (MGI Tech, Co., Ltd., China) and sequenced them on DNBSEQ-T7RS (MGI Tech, Co., Ltd.), generating 150-bp paired-end reads.

### RT-qPCR

Total RNA was extracted as described above. RT-qPCR was performed using ReverTra Ace qPCR RT Master Mix (Toyobo, Co., Ltd., Japan) and Fast SYBR Green Master Mix (Thermo Fisher Scientific, Inc.). Real-time PCR reactions were run on StepOne Real-Time PCR System (Thermo Fisher Scientific, Inc.). The six primer pairs shown in Supplementary Table 3 were tested to confirm their amplification efficiency. Based on the results, we selected PvTubulin_F1 and PvTubulin_R1 for amplifying PVIT_0011115 (*tubulin* of *P. viticola*) (Cruz-Silva et al. 2021) and VvEFalpha_F1 and VvEFalpha_R1 for amplifying CK203_026004 (*elongation factor 1-α* of *V. vinifera*) (Reid et al. 2006) to analyze the pathogen/host biomass ratio.

### Alignment of RNA-seq reads to reference genomes

RNA-seq reads from *P. viticola-*infected samples were first quality-filtered using fastp v0.23.2 with the default settings (Chen et al. 2018). Then, the filtered reads were aligned to the *P. viticola* isolate INRA-PV221 reference genome obtained from Recherche Data Gouv (https://entrepot.recherche.data.gouv.fr/dataset.xhtml?persistentId=doi:10.15454/4NYHD6) (Dussert et al. 2019) using STAR 2.7.10b with the --alignIntronMax 5000 option (Dobin et al. 2013), considering the maximum intron length in the *P. viticola* gene annotation (3,288 bp). The same filtered RNA-seq reads were also aligned to the *V. vinifera* reference genome (NCBI GenBank GCA_000003745.2) (Jaillon et al. 2007) using STAR 2.7.10b with the default settings.

### Heterozygous allele frequency calculation

The heterozygous allele frequency of *P. viticola* was computed using the method described by Bueno-Sancho et al. 2017 with minor modifications. The sequencing data used for this analysis are shown in Supplementary Tables 1, 4, and 5. Whole-genome sequencing (WGS) datasets were quality-filtered using fastp v0.23.2 with the default settings and aligned to the same reference genome using BWA-0.7.18 (r1243) with the default settings (Li 2013). RNA-seq datasets were quality-filtered and aligned to the reference genome as described above. The resulting alignment .sam files were converted into .bam files using samtools-1.16.1 with the view option (Danecek et al. 2021). SNPs were identified using samtools-1.16.1 with the mpileup option, applying a minimum depth of coverage of 10× for WGS datasets and 20× for RNA-seq datasets. At each SNP site, allele frequencies were assessed and those ranging from 0.2 to 0.8 were categorized as heterozygous sites and visualized using the R package ggplot2 4.0.2 (Wickham 2016).

### Identification of *tubulin* haplotype sequences

The .bam files of 14 peel sample RNA-seq datasets and 188 publicly available WGS datasets were generated as described above. These were used as input files to identify the *tubulin* haplotype sequences using samtools-1.23 with the parameters: consensus --mode simple --ambig --call-fract 0.2 --het-fract 0.2 --min-BQ 20 --min-MQ 20. For the 14 RNA-seq datasets, we investigated the average mapping depth of the *tubulin* coding region using samtools-1.23 with the depth -a option. Sequences containing heterozygous alleles were unfolded using DnaSP v6 (Rozas et al. 2017). The sequences were reconstructed into estimated haplotypes using the Excoffier-Laval-Balding (ELB) algorithm implemented in Arlequin ver 3.5.2.2 (Excoffier and Lischer 2010). The input file for running Arlequin was manually prepared based on a .tab file generated using SeqKit v2.11.0 with the fx2tab option (Shen et al. 2024). The identified *tubulin* haplotypes were summarized based on *tubulin* haplotype sequences documented by Fontaine et al. (2021) (https://entrepot.recherche.data.gouv.fr/dataset.xhtml?persistentId=doi:10.15454/FD86M2), using VSEARCH v2.30.6 with the --derep_fulllength option (Rognes et al. 2016). The maximum-likelihood tree of *tubulin* haplotypes was generated using RAxML-NG v1.2.2 with the -all option (Kozlov et al. 2019) based on alignments obtained using MUSCLE version 5.3 with the -align and -nt options (Edgar 2022). The alignments were manually inspected and visualized using MEGA 12 version 12.1.2 (Stecher et al. 2026) and CLC Genomics Workbench version 8.5.4 (QIAGEN N.V.). The phylogenetic tree was visualized using iTOL version 7.5 (Letunic and Bork 2024).

### Phylogenetic analysis based on the SNP profiles

We generated a phylogenetic tree using a total of 202 RNA-seq and WGS datasets, as illustrated in Supplementary Figure 1. The sequence data used for the analysis are described in Supplementary Tables 1 and 4. RNA-seq and WGS datasets were quality-filtered and aligned to the *P. viticola* reference genome as described above. The average mapping depth was determined using samtools-1.16.1 with the depth -a option to confirm that each WGS dataset had ≥10× average depth against the reference genome (Supplementary Table 6). SNP calling was performed using Freebayes 1.3.6 with the -- ploidy 2 --use-best-n-alleles 6 options (Garrison and Marth 2012). The raw variant calls in .vcf format were filtered as follows: First, the .vcf file was normalized using bcftools-1.17 with the norm -m -any options (Danecek et al. 2021). Quality filtering was then applied using vcffilter of VCFlib v1.0.15 with the parameter: -f “QUAL > 20 & QUAL / AO > 10 & SAF > 0 & SAR > 0 & RPR > 1 & RPL > 1 & AC > 0” (Garrison et al. 2022), as described by Sperschneider et al. (2023). Valid SNP sites were defined as those with a read depth ≥10× in at least 80% of the WGS datasets, and a read depth ≥ 20× in at least 80% of the RNA-seq datasets using bcftools with the view option. The extracted valid SNP sites were further filtered to keep only biallelic SNPs satisfying ≤ 20% sample missing rates and ≥ 5% minor allele frequency (MAF) using bcftools with the view option. The resulting .vcf file was converted to multiple sequence alignments in PHYLIP format using vcf2phylip v.2.9 (Ortiz 2019). The maximum-likelihood phylogenetic analysis using 13,275 SNP sites was conducted using RAxML-NG v1.2.2 with the --model TVM+G4+ASC_LEWIS option. The best-fit nucleotide substitution model was selected using ModelTest-NG v0.1.7 based on the Bayesian Information Criterion (BIC) (Darriba et al. 2020). The ASC_LEWIS option was used to correct for ascertainment bias since the input file does not contain constant sites (only SNPs). Bootstrap values were calculated using 1,000 replicates. The phylogenetic tree was visualized using iTOL version 7.5 (Letunic and Bork 2024).

We also generated a phylogenetic tree using only the 188 WGS datasets. The SNP calling using WGS datasets was independently performed using Freebayes 1.3.6 with the same options as above. The raw variant calls were also filtered in the same way, except for removing the thresholds for RNA-seq datasets. As the resulting .vcf file contains a very large number of SNP sites (1,333,776 sites), we extracted SNP sites at the third-codon positions of all CDSs in the reference genome annotation (145,015 sites) to reduce computational burden. The maximum-likelihood phylogenetic analysis using the SNPs at the third-codon positions was conducted as described above. Bootstrap values were calculated using 100 replicates. The phylogenetic tree was visualized using iTOL version 7.5 (Letunic and Bork 2024).

## Results

### Previously reported epiphytic tissue sampling methods are not applicable to *P. viticola*-infected grapevine leaves

We initially tested whether known epiphytic tissue sampling methods using cellulose acetate could be applied to *P. viticola*-infected leaves. Previously reported methods can be classified based on how the cellulose acetate solution is applied: immersing a leaf into the solution (Both et al. 2005), spreading the solution onto a leaf using a paintbrush (Kleemann et al. 2012), or dropping the solution onto a leaf using a pipette (McMullan et al. 2025). We first immersed 12-mm-diameter leaf discs or hand-cut leaf strips into the cellulose acetate solution. The immersed leaves were immediately retrieved and left for a few minutes to allow the acetone to evaporate. We found that the cellulose acetate film developed by this method was too thin and fragile to be removed from the leaf surface using forceps (Supplementary Figure 2A–C). Similarly, cellulose acetate applied with a paintbrush readily collapsed when we attempted to remove it from the leaf surface (Supplementary Figure 2D). Next, we dropped approximately 200 µL of the cellulose acetate solution onto lesion areas using a pipette. The resulting cellulose acetate film could be removed from the leaf surface using forceps (Supplementary Figure 2E, F). However, we found that it was difficult to retain the solution on the lesion areas because *V. vinifera* leaves are not completely flat, especially around the leaf veins. Additionally, evaporating acetone took longer (> 10 min) than the other two methods, because we applied an excessive amount of the cellulose acetate solution to obtain a thicker film to peel. Taken together, we concluded that the three previous methods are not suitable for *P. viticola*-infected grapevine leaves.

### Establishing a new epiphytic tissue sampling method using a cellulose acetate membrane filter

To overcome the shortcomings of the three methods tested, we developed a new sampling method using a cellulose acetate membrane filter. This method was inspired by a peel technique for preparing sections of fossil plants using a cellulose acetate film and acetone (Joy et al. 1956). Instead of the 0.05-mm-thick cellulose acetate film used in the original method, we used a 13-mm-diameter precut cellulose acetate membrane filter, which sufficiently covers a *P. viticola* lesion (Fig. 1A, B; Supplementary Video 1). Approximately 100 µl of acetone was added to a cellulose acetate membrane filter, partially dissolving the filter surface, which was then placed on a lesion (Fig. 1C; Supplementary Video 1). After a few minutes, the fixed cellulose acetate membrane filter was removed using forceps and collected in a 2.0-mL tube (Fig. 1D, E; Supplementary Video 1). Microscopic observation confirmed that the removed filter contained many sporangia and sporangiospores of *P. viticola* (Fig. 1F, G). Hereafter, we refer to this cellulose acetate membrane filter removed from the leaf surface as a “peel sample”.

**Figure 1.**
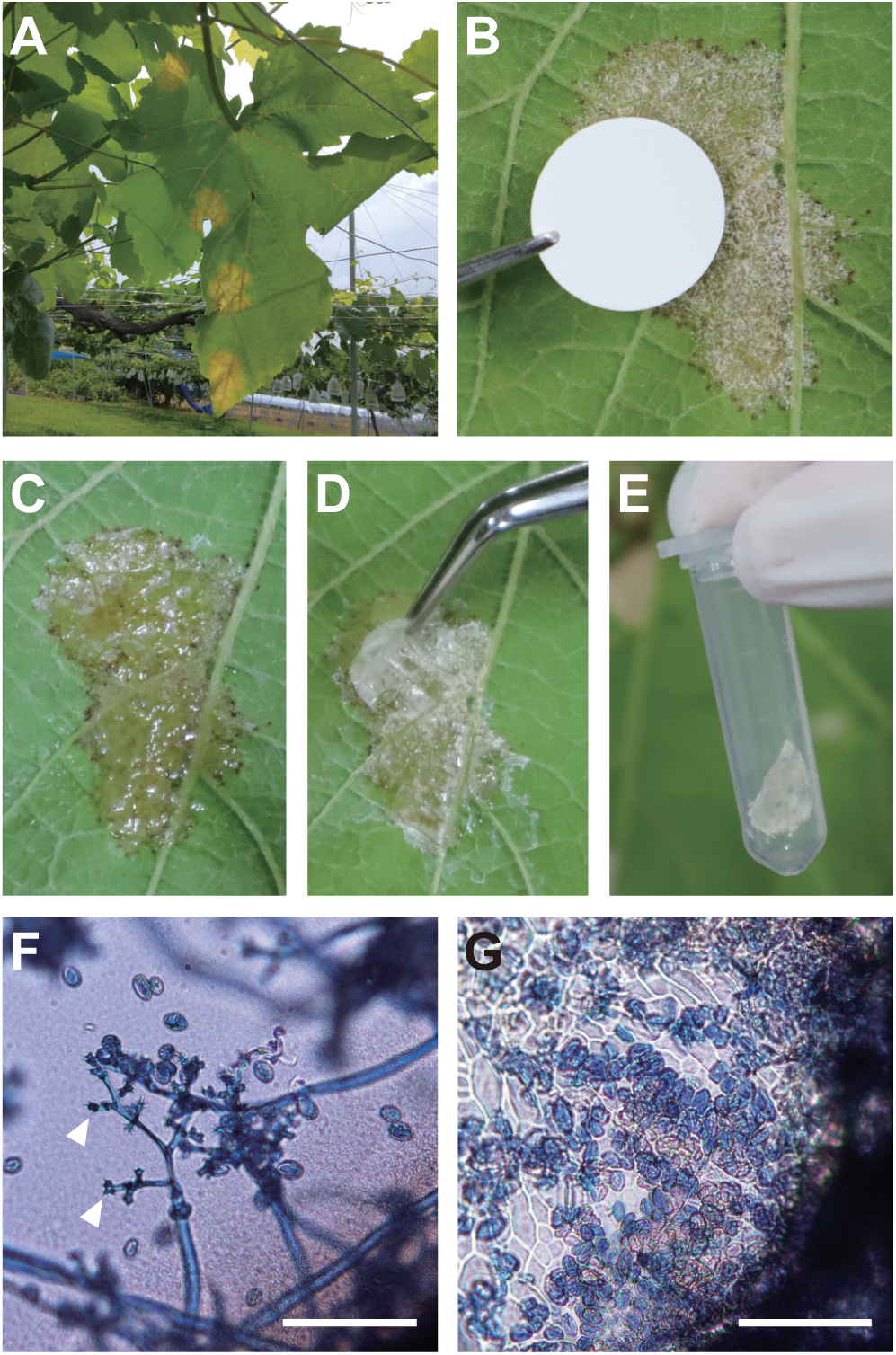
Epiphytic tissue sampling of a *P. viticola*-infected grapevine leaf using a cellulose acetate membrane filter. (A) A grapevine leaf infected by *P. viticola*. (B) A cellulose acetate membrane filter was placed on the abaxial leaf surface where *P. viticola* sporulates. (C) The cellulose acetate membrane filter was fixed on the leaf surface as acetone was evaporated. (D) The fixed membrane filter was removed from the leaf surface using forceps. (E) The removed membrane filter was collected in a sterilized 2.0-mL tube. (F–G) Microscopic observation of peel samples stained with trypan blue to visualize *P. viticola* sporangia (F, G) and sporangiospores (F, arrowheads). Scale bars = 100 µm.

### High-quality RNA was extracted from peel samples of *P. viticola*-infected leaves

We next tested whether high-quality RNA could be extracted from peel samples using a commercially available RNA extraction kit. We collected *P. viticola*-infected leaves from seven different grapevine cultivars and *Vitis ficifolia* var. *ganebu* in Osaka and Yamanashi, Japan, in 2022 and 2023. We then extracted total RNA from 20 leaf discs and 28 peel samples prepared from these infected leaves. The average RNA concentration assessed by NanoDrop was 19.8±24.6 ng/µl (mean±SD) in leaf disc samples and 64.9±43.6 ng/µl (mean±SD) in peel samples (Fig. 2A; Supplementary Table 2). The average A_260_/A_280_ ratio assessed by NanoDrop was 1.20±0.32 (mean±SD) in leaf disc samples and 1.89±0.13 (mean±SD) in peel samples (Fig. 2B; Supplementary Table 2). These results indicate that, in general, peel samples yield RNA of higher quantity and quality compared to leaf discs. Overall, we found that peel samples are more amenable to RNA extraction using a standard protocol than leaf discs.

**Figure 2.**
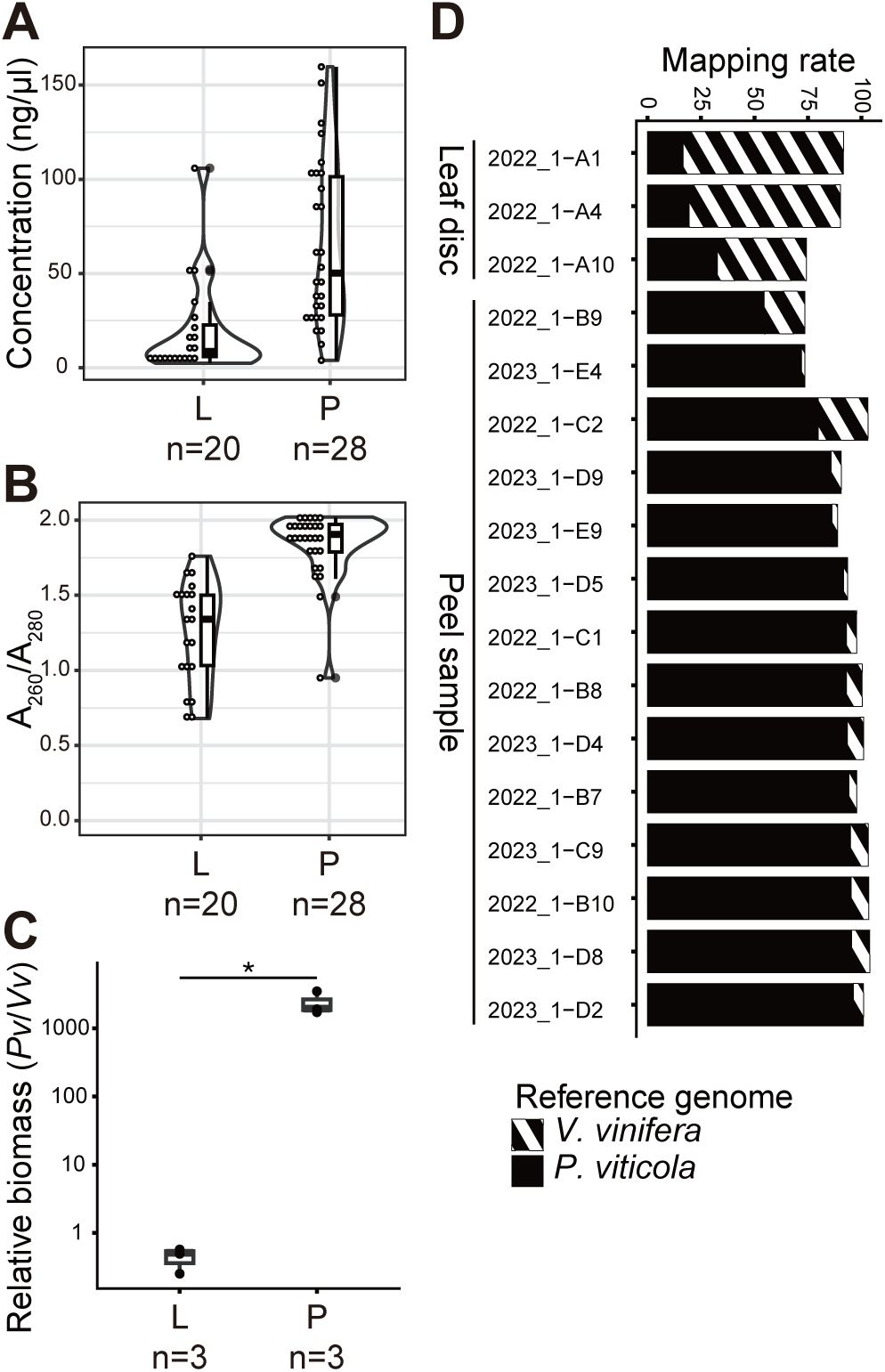
Peel samples yield higher RNA quantity and quality and enrich pathogen biomass more effectively than conventional leaf discs. (A) RNA concentration (ng/µL) measured by NanoDrop. (B) Ratio of absorbance values at 260 nm and 280 nm (A_260_/A_280_) of RNA measured by NanoDrop. (C) Relative pathogen/host biomass determined by RT-qPCR comparing the transcript abundances of *P. viticola tubulin* and *V. vinifera elongation factor 1-α*. The asterisk represents a significant difference (*P* < 0.05, two-tailed Student’s t-test). (D) Mapping rates of RNA-seq reads to the *P. viticola* and *V. vinifera* reference genomes. L and P indicate leaf discs and peel samples, respectively. N indicates the number of samples tested.

### Peel samples enrich *P. viticola* RNA more effectively than the conventional leaf disc samples

To assess whether peel samples enrich *P. viticola* biomass, we conducted RT-qPCR. This analysis showed that the relative *P. viticola* biomass in peel samples was more than 5,000-fold higher than that in leaf discs on average (Fig. 2C), confirming that pathogen biomass is enriched in peel samples compared to leaf discs. We then subjected three leaf disc samples and 14 peel samples to RNA-seq. The obtained RNA-seq reads were independently aligned to the *P. viticola* and *V. vinifera* reference genomes. The average mapping rate to the *P. viticola* reference genome was 22.9±8.6 % (mean±SD) for leaf disc samples and 87.5±11.8% (mean±SD) for peel samples (Fig. 2D). The average mapping rate to the *V. vinifera* reference genome was 62.5±18.2% (mean±SD) for leaf disc samples and 7.6±6.3% (mean±SD) for peel samples (Fig. 2D). These results indicate that the newly established sampling method effectively captures pathogen epiphytic biomass, leading to higher pathogen-derived RNA-seq read proportions in peel samples than in leaf discs.

### *P. viticola* in a single lesion of a field-collected leaf represents a predominantly single genotype

To characterize pathogen genotypes in field-collected samples without single spore isolation, a pathogen sample must represent a single genotype without cross-contamination. To address whether each field RNA-seq dataset was derived from a single genotype, we calculated the heterozygous allele frequency. Given that *P. viticola* propagates as a heterothallic diploid (Dussert et al. 2020), the distribution of heterozygous single-nucleotide polymorphism (SNP) site counts is expected to display a single mode at 0.5. We first analyzed publicly available whole-genome sequencing (WGS) datasets of seven *P. viticola* isolates obtained through single-sporangium isolation (Dussert et al. 2019, 2020). The heterozygous allele frequency histograms showed single modes at 0.5, as expected, indicating that these *P. viticola* isolates are diploid (Supplementary Figure 3A). We then analyzed publicly available RNA-seq datasets of two isolates obtained via single-sporangium isolation (Brilli et al. 2018; Dussert et al. 2019). The resulting histogram shapes were comparable to the WGS datasets, suggesting that *P. viticola* has little allele-specific expression bias (Supplementary Figure 3B). We thus conclude that RNA-seq data can be used to examine the *P. viticola* genotype purity based on the distribution of heterozygous alleles. Next, we analyzed our field RNA-seq datasets. The average number of sites with > 20× depth of coverage was 4.4×10^6^±2.9×10^6^ (mean ± SD) for leaf discs and 1.7×10^7^±4.2×10^6^ (mean ± SD) for peel samples, demonstrating that enriched pathogen reads in RNA-seq data increased the number of sites considered for detecting SNPs (Fig. 3A). Only two alleles were identified across all field-collected samples, with frequencies of a third allele being nearly zero (Fig. 3B, C). Except for the leaf disc sample “2022_1-A4”, which did not show a clear peak (Fig. 3B), heterozygous allele frequency histograms for all datasets exhibited single modes at 0.5, consistent with publicly available isolates originating from single spores. These findings suggest that *P. viticola* in a single lesion under field conditions predominantly represents a single genotype.

**Figure 3.**
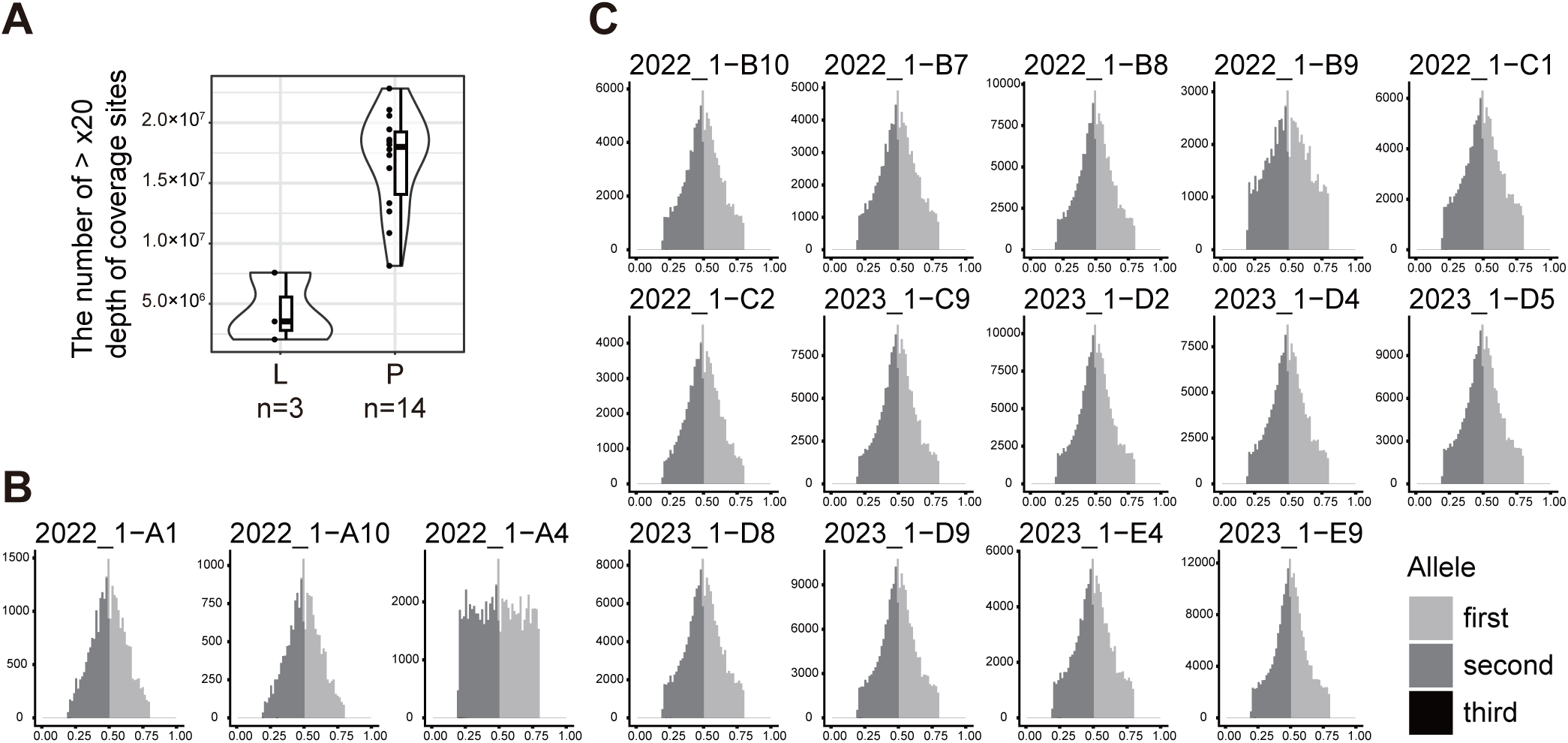
A *P. viticola* field sample derived from a single lesion represents a predominantly single genotype. (A) Number of sites with > 20× depth of coverage detected using RNA-seq reads mapped to the *P. viticola* reference genome. L and P indicate leaf discs and peel samples, respectively. N indicates the number of samples tested. (B, C) *P. viticola* field samples displayed a single mode at 0.5 for the frequency of heterozygous sites, which is typical for a single genotype in a diploid organism. The distribution of heterozygous site counts was determined for three leaf disc samples (B) and 14 peel samples (C). The x-axis represents the allele frequency, with values between 0.2 and 0.8 categorized as heterozygous. The y-axis represents the number of detected heterozygous sites.

### Japanese *P. viticola* samples tested belong to *P. viticola* f. sp. *aestivalis*

Previous studies reported five cryptic species or *formae speciales* within native *P. viticola* that exhibit distinct host ranges among wild Vitaceae plants (Rouxel et al. 2013, 2014). To identify the cryptic species of our 14 peel samples, we analyzed the *tubulin* haplotypes, which are well associated with the cryptic species classification (Rouxel et al. 2013, 2014). Alongside the RNA-seq datasets, we sourced publicly available WGS datasets of 188 *P. viticola* isolates; 187 were collected across nine European countries (France, Switzerland, Germany, Italy, Hungary, Spain, Bulgaria, Czechia, and Georgia) and one from China (Supplementary Table 4). We independently aligned a total of 202 RNA-seq and WGS datasets to the *P. viticola* reference genome and reconstructed *tubulin* haplotype sequences. This analysis identified four haplotypes documented by Fontaine et al. (2021) (Hap_3, Hap_10, Hap_11, and Hap_12), as well as three novel haplotype sequences from three isolates with WGS data (Pv1419_1, Pv3512, and Pv3716) (Supplementary Table 7; Supplementary Figure 4). Phylogenetic analysis classified all seven haplotype sequences as *P. viticola* f. sp. *aestivalis* (Supplementary Figure 5A). Among them, Hap_10 and Hap_11 were the most frequently identified haplotypes, accounting 87.4% of detected haplotypes (Supplementary Figure 5B). All 202 samples/isolates tested had at least one haplotype of either Hap_10 or Hap_11 (Supplementary Table 7). This finding is consistent with a previous study showing that Hap_10 and Hap_11 are the most common haplotypes among invasive *P. viticola* f. sp. *aestivalis* lineages (Fontaine et al. 2021). The 14 Japanese samples harbored Hap_10 and/or Hap_11 at their *tubulin* locus: eight were Hap_10/Hap_10 homozygous, five were Hap_10/Hap_11 heterozygous, and one was Hap_11/Hap_11 homozygous (Supplementary Figure 5C). These results suggest that our Japanese samples and the isolates with publicly available WGS datasets share similar *tubulin* haplotypes and are classified as *P. viticola* f. sp. *aestivalis*.

### Japanese *P. viticola* samples are genetically distinct from isolates collected in other geographic areas

To investigate the genetic background of Japanese *P. viticola* in detail, we conducted phylogenetic analysis using the RNA-seq datasets from our 14 peel samples and the publicly available WGS datasets from 188 isolates. We conducted SNP calling against the *P. viticola* reference genome using a total of 202 RNA-seq and WGS datasets. We then generated a maximum-likelihood phylogenetic tree with the 13,275 SNP sites that were detected in ≥ 80% of the 14 RNA-seq datasets to avoid overrepresenting SNPs identified in the WGS datasets only (Fig. 4). In the resulting tree, French isolates and isolates from other European countries and China tended to separate, although the basal nodes containing these isolates had low bootstrap support values (< 80%). We observed a similar tree topology with low bootstrap support values when analyzing 145,015 SNP sites detected across WGS datasets of 188 isolates (Supplementary Figure 6). These results imply that RNA-seq datasets can reconstruct phylogenies at a similar resolution to WGS datasets. The lower bootstrap support values reflect the documented genetic proximity among introduced *P. viticola* lineages (Fontaine et al. 2021). The 14 Japanese samples formed a single clade supported by a 100% bootstrap support value regardless of differences in collection years, sites, or host species/cultivars (Fig. 4). Interestingly, they were separated on the tree from an isolate collected in China in 2015 (YL), despite their geographic proximity. These findings suggest that our Japanese samples are genetically distinct from the European and Chinese isolates tested, even though they belong to the same *P. viticola* f. sp. *aestivalis* lineage.

**Figure 4.**
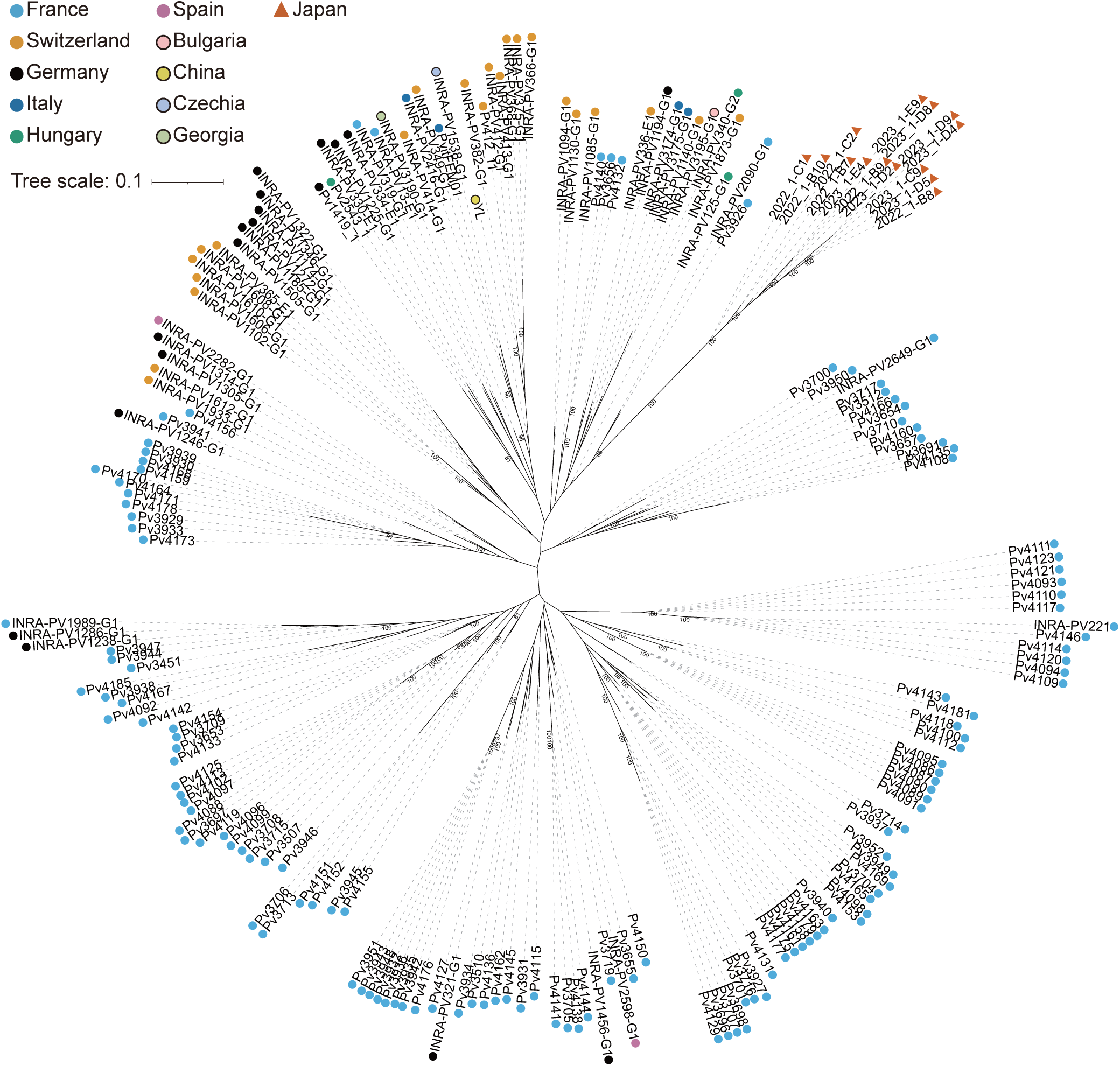
Japanese *P. viticola* samples are genetically distinct from isolates mainly collected from Europe. The maximum-likelihood tree was generated using 14 RNA-seq datasets of field samples and 188 publicly available WGS datasets of isolates, based on 13,275 SNP sites. For each sample/isolate, its country of origin is shown at the tip. Bootstrap values ≥ 80% are shown at the nodes. Triangles and squares indicate RNA-seq and WGS datasets, respectively.

## Discussion

Responding to invasive pathogens requires rapid and accurate surveillance and diagnostic systems. Here, we developed a sampling method that effectively collects epiphytic *P. viticola* material. Our analysis showed that the pathogen read proportion in peel samples increased by about threefold on average compared to those in conventional leaf discs. We hypothesize two reasons for this dramatic increase in pathogen biomass in peel samples. The first reason is that a cellulose acetate membrane filter effectively captured *P. viticola* on the *V. vinifera* leaf surface. Previously reported sampling methods using a cellulose acetate solution were used for pathogen-infected leaves of *Hordeum vulgare*, *Arabidopsis thaliana*, or *Beta vulgaris*, which have relatively smooth leaf surfaces (Both et al. 2005; Kleemann et al. 2012; McMullan et al. 2025). In contrast, the abaxial leaf surface of *V. vinifera*, where *P. viticola* sporulates, has highly elongated trichomes (Ma et al. 2016). Although this “hair coat” prevents a cellulose acetate solution from forming a readily removable strip, the preformed membrane assists in developing a stable strip on the complex *V. vinifera* surface. The second reason is the effective removal of host tissues containing metabolites that inhibit RNA extraction. We demonstrate that the total RNA yields extracted from *P. viticola* peel samples are higher than those extracted from conventional leaf discs. This is likely because our sampling method effectively eliminated grapevine tissues containing secondary metabolites that inhibit nucleic acid extraction. The new sampling method simplifies pathogen nucleic acid extraction from host plants rich in polysaccharides and polyphenols without requiring specialized protocols. Overall, the newly developed method enables high pathogen RNA yields from field samples infected by *P. viticola* that cannot be cultured on artificial media. In theory, this method can be applied to other pathogen-infected plant materials, particularly those with complex surface structures or metabolites that inhibit nucleic acid extraction.

Our results indicate that peel samples enrich more pathogen biomass than conventional leaf discs, allowing identification of more SNPs with high confidence. Since a higher-density SNP panel improves the accuracy of population genetic analyses in general (Georges-Filteau et al. 2020), our newly developed sampling method should aid genotyping and subsequent analyses of unculturable plant pathogens. Furthermore, our heterozygous allele frequency analysis shows that *P. viticola* colonizing a single lesion predominantly represents a single genotype in a field environment. This finding is similar to observations in *Puccinia striiformis* f. sp. *tritici* and *P. graminis* f. sp. *tritici*, where heterozygous allele distributions were investigated using field RNA-seq data (Hubbard et al. 2015; Tsushima et al. 2022). Based on our results, we propose that population genomic analyses using field NGS data, without single spore isolation, are also possible for *P. viticola*.

Our phylogenetic analysis suggests that the Japanese samples tested in this study are genetically distinct from isolates mainly collected in European countries. Interestingly, all the samples/isolates examined have similar *tubulin* haplotypes that are frequently identified among introduced *P. viticola* f. sp. *aestivalis* populations (Fontaine et al. 2021). These findings suggest substantial genetic variations even within the invasive *P. viticola* lineages. As the Japanese *P. viticola* population may possess unique genetic variations relevant to infection, such as avirulence/virulence effector genes, tailored control measures may be required. Historical records suggest that the Japanese *P. viticola* population was introduced from overseas (Kikuchi 1948; Nagasawa 1976). However, its genetic origin is uncertain, as the Japanese government at the time imported grapevine cultivars from multiple geographic areas, including France and the United States (Kikuchi 1948; Nagasawa 1976). Our work generates new working hypotheses regarding the immigration route of Japanese *P. viticola*. The first possible scenario is that Japanese *P. viticola* was introduced via Europe, as proposed by Fontaine et al. (2021), and obtained new genomic variations after introduction to Japan. The second possibility is that the Japanese *P. viticola* population was introduced from other geographic areas, for instance, North America, where native *P. viticola* f. sp. *aestivalis* populations are present. To test these hypotheses, detailed population genomic studies will be essential, using current and historical samples/isolates collected across Japan as well as other countries, especially the United States.

In conclusion, we have established a new method for effectively collecting field samples of *P. viticola*, an important obligate biotrophic pathogen in vineyards. Our phylogenetic analysis revealed that our Japanese samples collected with this method are genetically distinct from European and Chinese isolates examined. Obligate biotrophic plant pathogens, including *P. viticola,* are often understudied due to the technical challenges of culturing them in the laboratory. Our findings highlight the potential of this newly developed sampling technique to enhance pathogen monitoring by directly capturing their genetic snapshots from the field.

## Acknowledgements

We thank the Osaka Metropolitan University Education and Research Field, Masafumi Yamashige, Yuji Matsuo, and Atsushi Tsushima for sample collection. We also thank Kumiko Kitano and Maki Hazama for technical assistance, and Motoaki Tojo and Soichiro Asuke for useful discussion. This work was conducted in part using resources of the OMU Center for Shared Research Equipment at Osaka Metropolitan University. Bioinformatic analysis was partially performed on the NIG supercomputer at the Research Organization of Information and Systems (ROIS) National Institute of Genetics (NIG).

## Data availability

The sequencing data obtained in this study are deposited in the NCBI Sequence Read Archive (SRA) under the BioProject ID PRJNA1556380. The scripts used in this study are available in a GitHub repository (https://github.com/TsushimaLab/peel_technique). Some initial scripts for computational analyses were partially generated using Gemini and ChatGPT; the authors manually reviewed and validated them.

## List of supplementary files

**Supplementary Table 1.** RNA-seq datasets used in this study.

**Supplementary Table 2.** Concentration and quality of extracted RNA.

**Supplementary Table 3.** Primers used for RT-qPCR.

**Supplementary Table 4.** Publicly available genome sequencing datasets used in this study.

**Supplementary Table 5.** Publicly available RNA-seq datasets used in this study.

**Supplementary Table 6.** Average mapping depth of WGS datasets.

**Supplementary Table 7.** *tubulin* haplotype sequences detected in this study.

**Supplementary Figure 1.** Schematic illustrating the steps of maximum-likelihood analysis based on SNPs. A total of 202 datasets were independently mapped to the reference genome. After SNP calling, detected SNPs were filtered based on quality and read depth. Biallelic SNPs with ≤ 20% sample missing rates and ≥ 5% minor allele frequency were used for maximum-likelihood (ML) tree construction.

**Supplementary Figure 2.** Testing previously reported sampling methods using cellulose acetate solution. (A, B) Hand-cut leaf strips (A) and 12-mm-diameter leaf discs (B) immersed in cellulose acetate solution and allowed to evaporate acetone on aluminum foil. (C) Cellulose acetate films developed on leaf discs immersed in cellulose acetate solution were fragile and difficult to remove. (D) Application of cellulose acetate solution onto the leaf surface using a paintbrush. (E) Dropping approximately 200 µL of cellulose acetate solution onto the leaf surface using a pipette. (F) Removed films formed by dropping cellulose acetate solution.

**Supplementary Figure 3.** (A) Frequency of heterozygous sites detected in WGS datasets of seven *P. viticola* isolates published by Dussert et al. (2019, 2020). (B) Frequency of heterozygous sites detected in RNA-seq datasets of two *P. viticola* isolates published by Brilli et al. (2018) and Dussert et al. (2019). RNA-seq datasets collected at different hours post-inoculation (hpi) were examined for each isolate. The x-axis represents the allele frequency, with values between 0.2 and 0.8 categorized as heterozygous. The y-axis represents the number of detected heterozygous sites. Note that both WGS and RNA-seq datasets were analyzed for the isolate INRA-PV221.

**Supplementary Figure 4.** Alignments of the seven *tubulin* haplotype nucleotide sequences detected across the 202 datasets. Dots indicate identical sequences to Hap_10, the most frequently detected haplotype. Variable sequence positions are highlighted with a pink background.

**Supplementary Figure 5.** (A) Maximum-likelihood tree constructed using the *tubulin* haplotypes documented by Fontaine et al. (2021) and the three newly identified haplotypes from three *P. viticola* isolates (Pv1419_1, Pv3512, and Pv3716). The *tubulin* sequence of *Plasmopara halstedii* was used as an outgroup. Bootstrap values ≥ 80% are shown at the nodes. (B) Frequency of *tubulin* haplotypes detected across the 202 datasets. Each dataset contributes two haplotype counts because *P. viticola* is diploid. Two counts were scored for a haplotype in a homozygote, while one count was scored for each haplotype in a heterozygote. (C) Frequency of *tubulin* haplotype combinations among the 14 peel samples collected in Japan.

**Supplementary Figure 6.** Maximum-likelihood tree generated using 188 publicly available WGS datasets of isolates, based on 145,015 SNP sites at the third codon positions. For each isolate, its country of origin is shown on the tip. Bootstrap values ≥ 80% are shown at the nodes.

**Supplementary Video 1.** Sampling procedure for the grapevine epiphytic tissue infected by *P. viticola* using a cellulose acetate membrane filter.

## Author contributions

A.T. conceived the project; C.K. and A.T. prepared figures and supplementary files; A.T., C.K., and K.A. wrote the manuscript; C.K., A.T., and S.S. collected field samples; C.K. extracted RNA from field samples and investigated mapping rates and allele frequencies; A.T. designed the sampling method, conducted microscopic analysis and RT-qPCR, and analyzed *tubulin* haplotype sequences; C.K. and A.T. performed SNP calling and phylogenetic analysis; All authors have read and approved the final manuscript.

## Declarations for conflict of interest and ethical standards

The authors declare no conflicts of interest or ethical concerns.

