## Supplementary Figure for "An epiphytic tissue sampling method using a cellulose acetate membrane filter facilitates phylogenomic analysis of *Plasmopara viticola*"

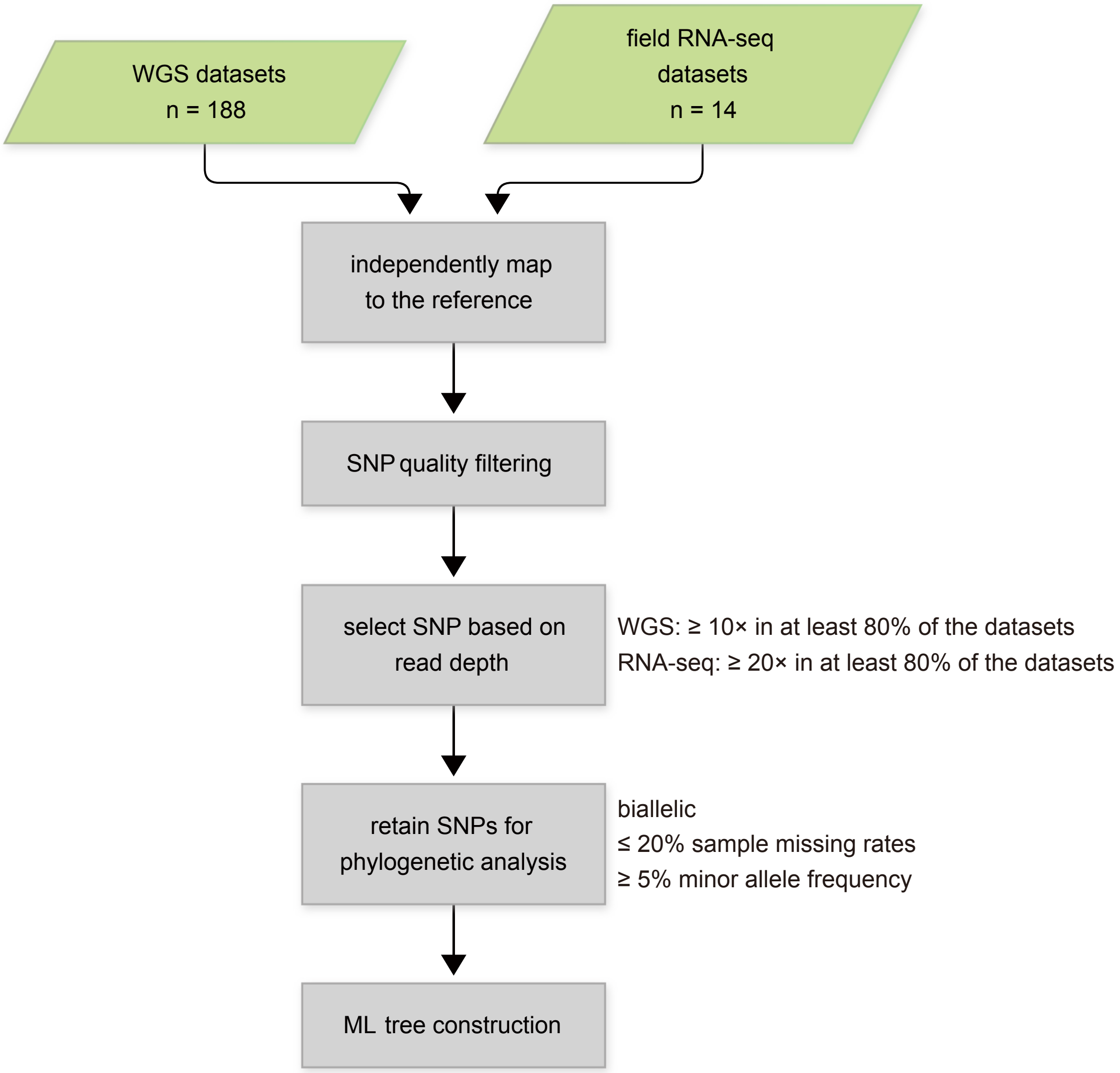


**Supplementary Figure 1.** Schematic illustrating the steps of maximum-likelihood analysis based on SNPs. A total of 202 datasets were independently mapped to the reference genome. After SNP calling, detected SNPs were filtered based on quality and read depth. Biallelic SNPs with ≤ 20% sample missing rates and ≥ 5% minor allele frequency were used for maximum-likelihood (ML) tree construction.

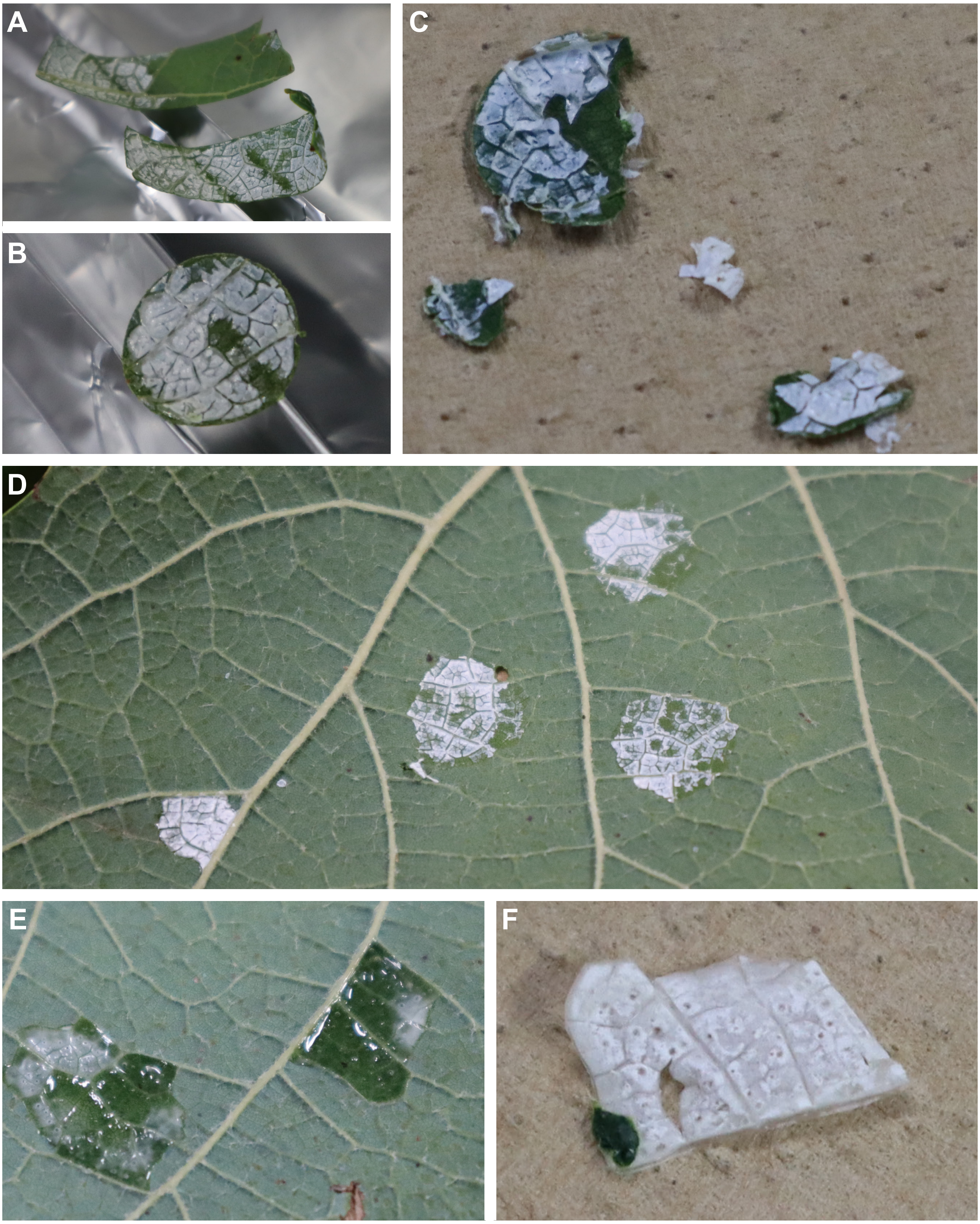


**Supplementary Figure 2.** Testing previously reported sampling methods using cellulose acetate solution. (A, B) Hand-cut leaf strips (A) and 12-mm-diameter leaf discs (B) immersed in cellulose acetate solution and allowed to evaporate acetone on aluminum foil. (C) Cellulose acetate films developed on leaf discs immersed in cellulose acetate solution were fragile and difficult to remove. (D) Application of cellulose acetate solution onto the leaf surface using a paintbrush. (E) Dropping approximately 200 µL of cellulose acetate solution onto the leaf surface using a pipette. (F) Removed films formed by dropping cellulose acetate solution.

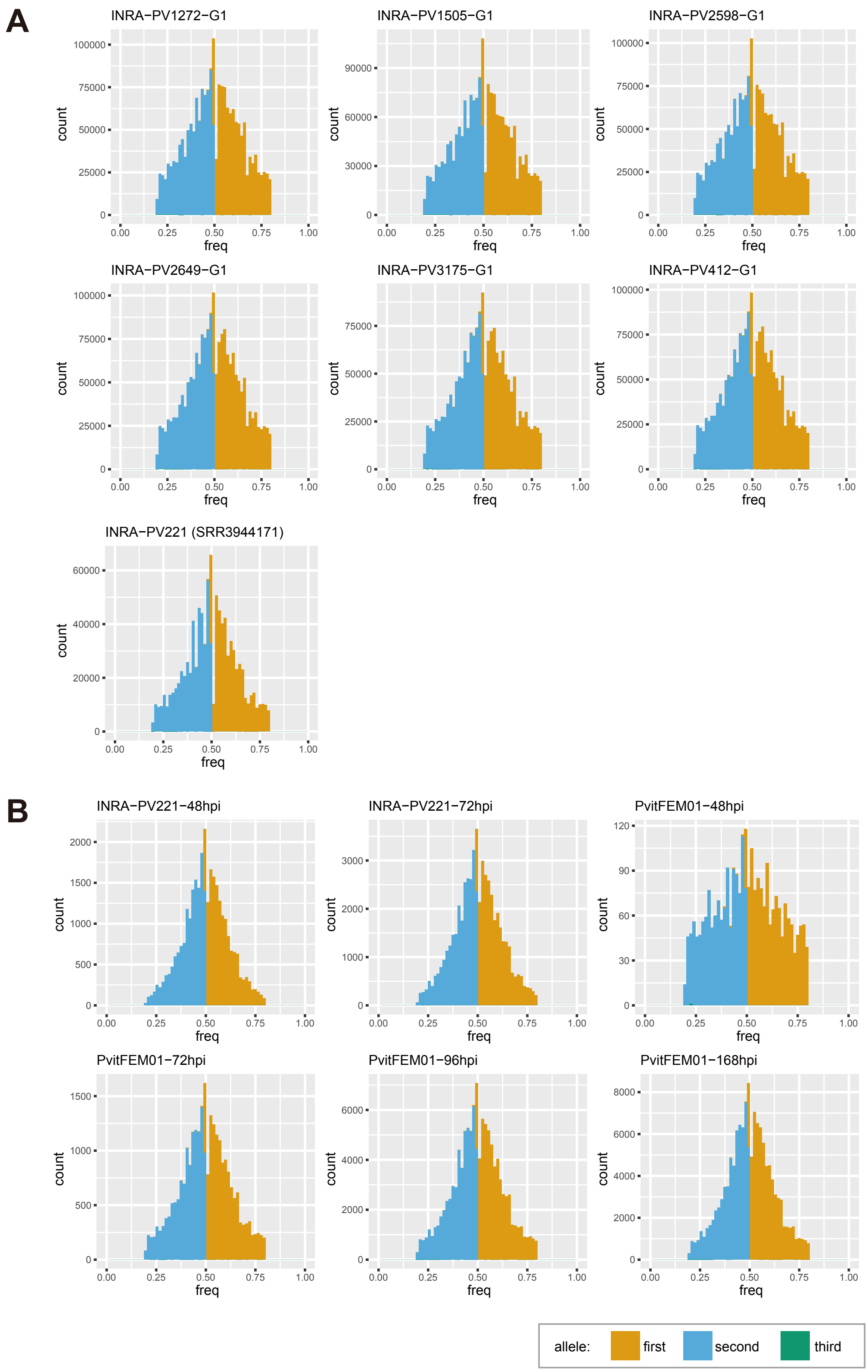


**Supplementary Figure 3.** (A) Frequency of heterozygous sites detected in WGS datasets of seven *P. viticola* isolates published by Dussert et al. (2019, 2020). (B) Frequency of heterozygous sites detected in RNA-seq datasets of two *P. viticola* isolates published by Brilli et al. (2018) and Dussert et al. (2019). RNA-seq datasets collected at different hours post-inoculation (hpi) were examined for each isolate. The x-axis represents the allele frequency, with values between 0.2 and 0.8 categorized as heterozygous. The y-axis represents the number of detected heterozygous sites. Note that both WGS and RNA-seq datasets were analyzed for the isolate INRA-PV221.

```
                                          20                            40                            60                            80
                                           |                             |                             |                             |
Hap_10    GACGAGGTCA TGTGCCTGGA CAATGAAGCC TTGTACGACA TTTGCTTTCG TACTCTGAAA CTCACCACTC CCACGTACGG 80
Hap_11    .......... .......... .......... .......... .......... .......... .......... .......... 80
Hap_12    .......... .......... .......... .......... .......... .......... .......... .......... 80
Hap_3     .......... .......... .......... .......... .......... .......... .......... .......... 80
Pv1419_1  .......... .....T.... T......... .......... .......... ......A... .......... .......... 80
Pv3512    .......... .......... .......... .......... .......... .......... .......... .......... 80
Pv3716    .......... .......... .......... .......... .......... .......... .......... .......... 80

                                         100                           120                           140                           160
                                           |                             |                             |                             |
Hap_10    CGACTTGAAT CACTTGGTTT GTGCCGCCAT GTCTGGCATC ACCACATGCC TTCGATTTCC CGGACAGCTG AATTCGGACT 160
Hap_11    .......... .......... .......... .......... .......... .......... .......... .......... 160
Hap_12    ....C..... .......... .......... .......... .......... .......... .......... .......... 160
Hap_3     .......... .......... .......... .......... .......... .......... .......... .......... 160
Pv1419_1  .......... .......... .......... .........T ..T....... .......... .......... .......... 160
Pv3512    ....C..... .......... .......... .......... .......... .......... .......... .......... 160
Pv3716    .......... .......... .......... .......... .......... .......... .......... .......... 160

                                         180                           200                           220                           240
                                           |                             |                             |                             |
Hap_10    TACGAAAATT GGCTGTTAAC CTGATTCCGT TCCCCCGTCT TCATTTCTTT ATGATTGGAT TTGCACCACT GACATCGCGC 240
Hap_11    .......... .......... .......... .......... .......... .......... .......... .......... 240
Hap_12    .......... .........T .......... .......... .......... .......... ........T. .......... 240
Hap_3     .......... .......... .......... .......... .......... .......... .......... .......... 240
Pv1419_1  .......... .......... .......... .......... .......... .......... .......... .......... 240
Pv3512    .......... .......... .......... .......... .......... .......... .......... .......... 240
Pv3716    .......... .......... .......... .......... .......... .......... .......... .......... 240

                                         260                           280                           300                           320
                                           |                             |                             |                             |
Hap_10    GGCTCGCAGC AGTATCGTGC GTTAACTGTA CCTGAGTTAA CCCAGCAGCA GTTTGATGCA AAAAATATGA TGTGTGCCGC 320
Hap_11    .......... .......... ...G...... .......... .......... .......... .......... .......... 320
Hap_12    .......... .......... ...G...... .......... .......... .......... .......... .......... 320
Hap_3     .......... .......... ...G...... .......... .......... .......... .......... .......... 320
Pv1419_1  .......... .......... ...G...... ........G. .......... .......... ..G....... ....C..... 320
Pv3512    .......... .......... .......... .......... .......... .......... .......... .......... 320
Pv3716    .......... .......... .......... .......... .......... .......... .......... .......... 320

                                         340                           360                           380                           400
                                           |                             |                             |                             |
Hap_10    TGATCCTCGC CACGGCCGCT ATTTAACAGC TGCGTGTATG TTCCGCGGAC GTATGAGCAC GAAGGAGGTT GACGAGCAGA 400
Hap_11    .......... .......... .......... .......... .......... .......... .......... .......... 400
Hap_12    .......... .......... .......... .......... .......... .......... .......... .......... 400
Hap_3     .......... .......... .......... .......... .......... .......... .......... .......... 400
Pv1419_1  .......... .......... .......... .......... .......... .C.....T.. .......... .......... 400
Pv3512    .......... .......... .......... .......... .......... .......... .......... .......... 400
Pv3716    .......... .......... .......... .......... .......... .......... .......... .......... 400

                                         420                           440                           460                           480
                                           |                             |                             |                             |
Hap_10    TGCTAAATGT TCAGAACAAA AACTCTTCGT ACTTTGTTGA ATGGATTCCC AACAATATCA AGGCTAGTGT GTGTGACATT 480
Hap_11    ....T..... .......... .......... .......... .......... .......... .......... .......... 480
Hap_12    .......... .......... .......... .......... .......... .......... .......... .......... 480
Hap_3     .......... .......... .......... .......... .......... .......... .......... .......... 480
Pv1419_1  .......... .......... .......... .......... .......... .......... .......... .......... 480
Pv3512    .......... .......... .......... .......... .......... .......... .......... .......... 480
Pv3716    ....T..... .......... .......... .......... .......... .......... .......... .......... 480

Hap_10    CCGCCCAAGG GCCTGAAAA 499
Hap_11    .......... ......... 499
Hap_12    .......... ......... 499
Hap_3     .......... ......... 499
Pv1419_1  ........A. ......... 499
Pv3512    .......... ......... 499
Pv3716    .......... ......... 499
```

**Supplementary Figure 4.** Alignments of the seven *tubulin* haplotype nucleotide sequences detected across the 202 datasets. Dots indicate identical sequences to Hap_10, the most frequently detected haplotype. Variable sequence positions are highlighted with a pink background.

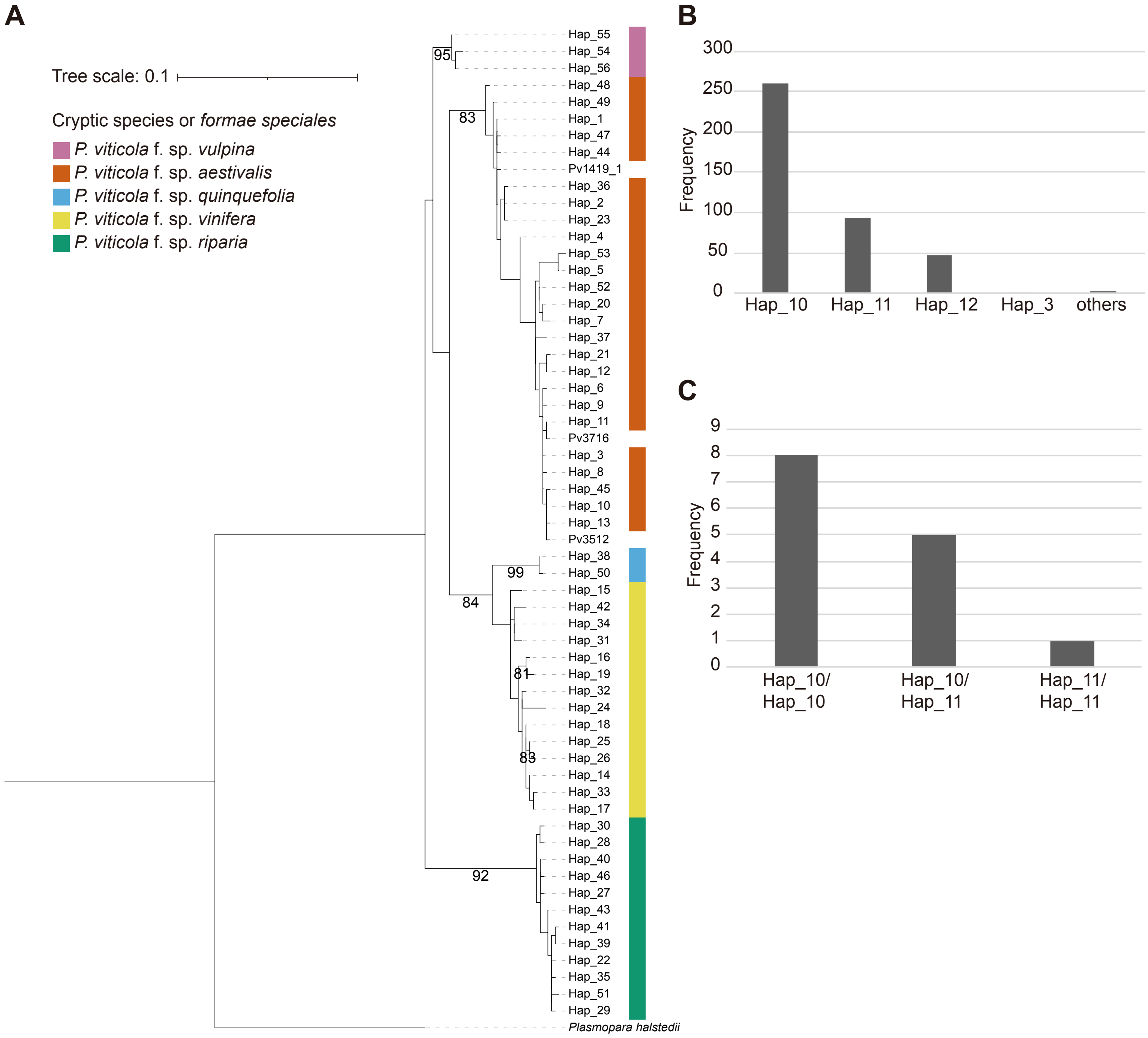


**Supplementary Figure 5.** (A) Maximum-likelihood tree constructed using the *tubulin* haplotypes documented by Fontaine et al. (2021) and the three newly identified haplotypes from three *P. viticola* isolates (Pv1419_1, Pv3512, and Pv3716). The *tubulin* sequence of *Plasmopara halstedii* was used as an outgroup. Bootstrap values ≥ 80% are shown at the nodes. (B) Frequency of *tubulin* haplotypes detected across the 202 datasets. Each dataset contributes two haplotype counts because *P. viticola* is diploid. Two counts were scored for a haplotype in a homozygote, while one count was scored for each haplotype in a heterozygote. (C) Frequency of *tubulin* haplotype combinations among the 14 peel samples collected in Japan.

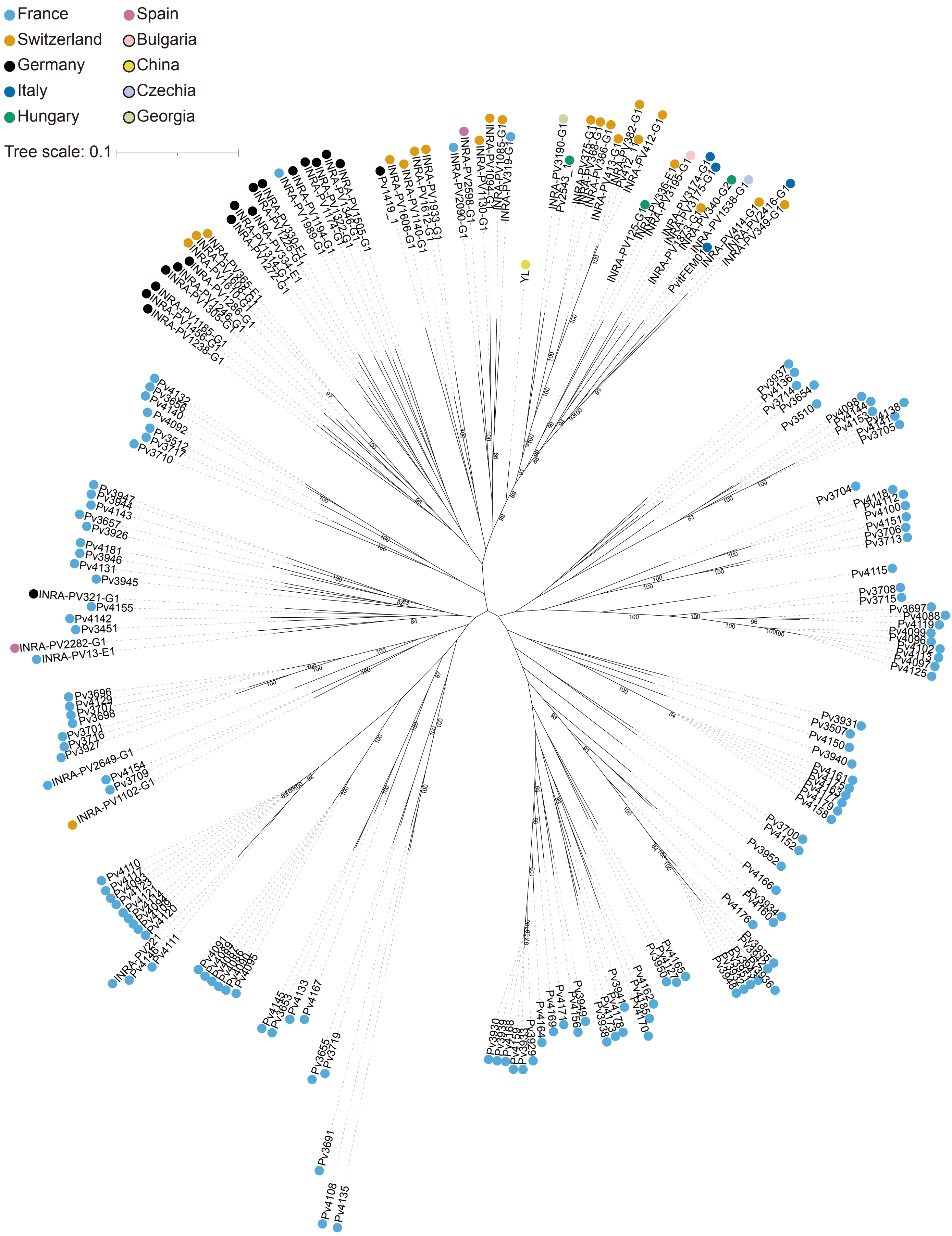


**Supplementary Figure 6.** Maximum-likelihood tree generated using 188 publicly available WGS datasets of isolates, based on 145,015 SNP sites at the third codon positions. For each isolate, its country of origin is shown on the tip. Bootstrap values ≥ 80% are shown at the nodes.
